# Visualizing TERRA RNA G-quadruplex unfolding in FUS biomolecular condensates

**DOI:** 10.1101/2025.10.29.685336

**Authors:** Tongyin Zheng, Nicolas L Fawzi

## Abstract

RNA G-quadruplexes (rG4s) are remarkably stable secondary structures with critical regulatory roles in gene expression, RNA metabolism, and telomere maintenance. However, their behavior within cells remains controversial, partly due to challenges in detecting rG4s in complex environments. Here, we use solution NMR spectroscopy to investigate how condensates formed by the low-complexity and RGG domains of the RNA-binding protein FUS affect the structure of TERRA, a highly stable model rG4 RNA. We show that FUS LC-RGG1 interacts with TERRA in dilute solution and that binding perturbs, but does not disrupt, the G-quadruplex structure. When co-phase separated with FUS LC-RGG1, however, NMR signatures of TERRA’s folded state disappear, and the remaining observable resonances indicate an unfolded conformation, even in buffer containing potassium where TERRA rG4 is exceptionally stable when outside a condensate. Quantitative comparisons with a mutant form of TERRA, used as a baseline for fully unfolded RNA, suggest that at minimum a third of TERRA RNA becomes unfolded in the condensed phase. Thus, our results demonstrate that condensates can shift the structural ensemble of rG4 towards unfolded species, offering a potential mechanistic explanation for their apparent lack of stability *in vivo* and revealing how phase-separated environments may actively modulate RNA structure and function.

**Highlights:**

- NMR spectroscopy directly probes RNA structure inside biomolecular condensate.
- FUS binding in solution perturbs TERRA RNA G-quadruplex.
- A sizable fraction of TERRA is unfolded in FUS condensates.

## Introduction

RNA G-quadruplexes (rG4s) are stable, four-stranded structures formed by guanine-rich RNA sequences that play essential regulatory roles in gene expression, RNA processing, and telomere maintenance (1–4). rG4 sequences are extremely stable in biochemical studies and maintain structure even when bound by proteins in molar excess (5) as well as in cell lysate conditions (6–8). Yet, the relevance and behavior of rG4s in cells remain controversial, as these same sequences that are stable in test tube conditions show evidence for lack of structure in cells by chemical probing approaches (3, 9). Why many G-quadruplexes show stability in test tubes and lack of evidence for stability in cells remains a question, in part due to challenges in reliably detecting and characterizing them in the complex cellular environment. One potential factor contributing to this mystery is the observation that G-quadruplex structures are often recruited to biomolecular condensates, membraneless cellular compartments formed through protein and nucleic acid phase separation (10–13). Various functions have been described for biomolecular condensate, such as dynamic reservoirs for proteins and RNAs, and hubs facilitating specific molecular interactions that differ significantly from those occurring in dilute solutions (14, 15). Specifically, G-quadruplexes have been proposed to participate in formation of condensates at transcription start sites, telomeres, nucleoli, nuclear speckles, and paraspeckles (16) and co-localize with phase separating RNA binding proteins in cells (17). Hence, it is possible that biomolecular condensates recruit G-quadruplex RNAs and their unique environment leads to unfolding of G-quadruplexes. Due to the highly concentrated nature of these environments, single-molecule approaches are often infeasible, and traditional biophysical methods, such as circular dichroism (CD) spectroscopy, are limited by strong background signals from proteins. These challenges suggest solution NMR may be a useful spectroscopic tool. Indeed, a recent study proposed that a modified G-quadruplex (MH24) from telomeres unfolds in condensates formed by a disordered domain of DDX4, an RNA helicase (16). However, several questions remain. This study used ^19^F NMR to probe the structure via a single fluorine-modified G base only and in dilute solution (buffer only) two resonances were observed, suggesting a conformational equilibrium as opposed to a single folded state. Addition of MH24 in condensates caused collapse of the ^19^F signals to a single broad resonance near that seen for thermally unfolded MH24 in dilute solution, however no structural details could be probed. Most importantly, rG4s are nearly universally stabilized by potassium ions placed at the core of the quadruplex yet the model condensates in the previous study were created in potassium-free buffer, potentially resulting in unfolding simply due to removal of the essential metal (16). Hence, it is important to directly probe the structural details of rG4 sequences that form well-established stable folds with high precision in physiologically relevant buffers.

The RNA-binding protein Fused in Sarcoma (FUS) is the archetype for phase separation via intrinsically disordered regions. FUS localizes to biomolecular condensates, and its biological function and disease-associated dysfunction are closely tied to its phase behavior (18, 19). Importantly, aberrant aggregation of FUS is associated with cellular toxicity in neurodegenerative diseases (20–22). However, recent studies have also suggested a protective role for FUS in C9orf72 ALS where GGGGCC repeat expansions (G4C2) are associated with cellular toxicity. Specifically, FUS has been proposed to function as a chaperone modulating the RNA G-quadruplex structures formed by G4C2 repeat expansions, potentially reducing their pathogenicity (23, 24). While this hypothesis is compelling, direct biophysical evidence for FUS-mediated rG4 destabilization is still lacking.

To investigate this proposed rG4-destabilizing function of FUS in a controlled biophysical context, we turned to a well-established model system. Telomeric repeat-containing RNA (TERRA) provides a robust platform for rG4 studies, thanks to its well-characterized and highly stable G-quadruplex structure (25, 26). In addition to its value as a model, TERRA plays essential roles in telomere biology and genome stability. In this study, we use solution NMR to probe the molecular details and structural consequences of the interaction between FUS and the highly studied dimeric G-quadruplex formed by TERRA12, a 12 nucleotide TERRA-derived RNA. Our goal is to provide mechanistic insight into how protein-RNA interactions within biomolecular condensates influence RNA structure and dynamics.

## Materials and Methods

### Oligonucleotides

RNA oligonucleotides were purchased from Horizon/Dharmacon and delivered HPLC-purified and lyophilized. TERRA12 (sequence UAGGGUUAGGGU) and a non-G4-forming TERRA mutant (27) (TERRA-mut; sequence UACCGUUACCGU) were resuspended in nuclease-free water to 1–2 mM. Folded TERRA RNAs were heated to 95 °C for 5 min in 20mM potassium phosphate pH 6.2, 50mM KCl buffer then slowly cooled to room temperature and equilibrated ~1 h at 25 °C. For unfolded controls, RNA was prepared in K^+^-free, 20mM MES pH 6.2 buffer. Poly(U) RNA (Sigma-Aldrich) was resuspended similarly and desalted with Zeba spin columns (Thermo, 7 kDa MWCO) into the indicated experimental buffer. All solutions and plastics were RNase-free.

### Protein expression and purification

Hexahistidine-tagged human FUS LC-RGG1 (residues 1–284) was expressed in *E. coli* BL21 Star (DE3) and purified from inclusion bodies as described previously (28) with the following details. Cells were grown at 37 °C in M9 minimal medium (27 mM NaCl, 22 mM KH_2_PO_4_, 51 mM Na_2_HPO_4_·7H_2_O, 1 mM MgSO_4_, 1% v/v MEM vitamins, trace metals “solution Q”, 4 g/L glucose, 1 g/L NH_4_Cl). For isotope labeling of FUS LC-RGG1, ^15^N-NH_4_Cl was used as the sole nitrogen source. At OD^600^ ~0.8, expression was induced with 1 mM IPTG (isopropyl β-D-1-thiogalactopyranoside) for 4 h at 37 °C. Cells were harvested (7,800 g, 15 min, 4 °C), frozen at −80 °C, then lysed in 20 mM sodium phosphate, 300 mM NaCl, pH 7.4 using a Fisherbrand Model 120 Sonic Dismembrator. Insoluble material was pelleted (47,855 g, 50 min, 4 °C), resuspended in the same buffer containing 8 M urea and 10 mM imidazole, clarified again, and filtered (0.2 µm). Ni^2+^-IMAC (5 mL HisTrap HP, Cytiva) was performed under denaturing conditions; bound protein was eluted with 300 mM imidazole in 20 mM sodium phosphate, 300 mM NaCl, 8 M urea, pH 7.4. Fractions were pooled and diluted ten-fold into 20 mM sodium phosphate, pH 7.4 to remove urea for TEV cleavage. His_6_-TEV protease 1:50-1:100 w/w protease:substrate was added and incubated overnight at room temperature. The reaction was returned to 8 M urea, 10 mM imidazole, and passed over the HisTrap again to remove the His tag and His-TEV. The flow-through containing tag-free FUS LC-RGG1 was buffer-exchanged into 20 mM CAPS, pH 11.0 (to deprotonate tyrosines and suppress phase separation), concentrated with 3 kDa MWCO Amicon filters, aliquoted, flash-frozen, and stored at −80 °C. Purity (>95%) was verified by SDS–PAGE. The “solution Q” trace-metal mix contained 8% v/v 5 M HCl, 0.5% w/v FeCl_2_·4H_2_O, 0.018% w/v CaCl_2_·2H_2_O, 0.0064% w/v H_3_BO_3_, 0.0018% w/v CoCl_2_·6H_2_O, 0.0004% w/v CuCl_2_·2H_2_O, 0.034% w/v ZnCl_2_, 0.0040% w/v Na_2_MoO_4_·2H_2_O.

### NMR Sample preparation

All NMR buffers were 20 mM potassium phosphate pH 6.2, 50 mM KCl unless noted otherwise, with 5% ^2^H_2_O except for condensed phase samples with 10% ^2^H_2_O (to improve lock signal). Dispersed-phase FUS-RNA samples were made with ^15^N-labeled FUS LC-RGG1 at 25 µM, mixed with TERRA12 at 110 µM (protein:RNA ≈ 1:4.4 molar) or with poly(U) at the same nucleotide concentration. Samples were optically clear with no visible condensates. Condensed-phase samples were made as described earlier (28). Specifically. macroscopic condensates were generated by rapid ten-fold dilution of concentrated LC-RGG1 stock (~3 mM in CAPS pH 11.0) into potassium phosphate buffer at pH 6.2 with 50 mM KCl and 10% ^2^H_2_O, followed by centrifugation at 4,000 g for 10 min at 25 °C. For RNA-containing condensates, RNA was added before centrifugation at a protein:RNA mass ratio of 20:1, corresponding to ~2.6:1 molar for TERRA12. After phase formation, the condensed material was transferred into a 3 mm NMR tube and compacted by brief centrifugations at 1,000 g. RNA incorporation into the condensed phase was quantified by measuring the A260 and A280 of the supernatant before and after condensation; incorporation was above 95% in all cases.

### NMR spectroscopy

NMR spectra were acquired at 298 K on a Bruker Avance III HD 850 MHz spectrometer with a TCI HCN cryoprobe and z-gradients. Data were processed in NMRPipe and analyzed in CCPNMR 2.5.2. <u>1D ^1^H</u>: water-suppressed ^1^H spectra were collected with the water resonance centered at 4.70 ppm, spectral width 16 ppm, TD = 32,768, and 16–128 scans. <u>2D ^1^H-^15^N HSQC</u>: 4,096 complex points (t_2_) × 400 increments (t_1_) with spectral widths of 10.5 ppm (^1^H) and 20 ppm (^15^N); carrier offsets 4.70 ppm (^1^H) and 117 ppm (^15^N). <u>2D ^1^H-^13^C HSQC of RNA</u> (natural abundance): 2D 1H-13C HSQCs were acquired with sensitivity-enhanced, gradient-selected HSQC. Two spectral windows were collected: 1) a C5–H5 window with 3,072 complex points in t_2_ and 200 increments in t_1_, spectral widths 12 ppm (^1^H) by 80 ppm (^13^C); and 2) an aromatic window for A C2–H2, U/C C6–H6, and A/G C8–H8 with 3,072 by 80 points, spectral widths 12 by 40 ppm. The ^1^H carrier was 4.70 ppm, and the ^13^C carrier was 80 ppm and 140 ppm, respectively. Assignments used literature chemical-shift regions for U C5–H5/C6–H6, A C8–H8/C2–H2, and G C8–H8.

### RNA electrophoresis

Denaturing polyacrylamide electrophoresis was used to assess RNA integrity. Samples containing RNA were mixed 1:1 with 2x denaturing loading buffer, heated to 70 °C for 3–5 min, and snap-cooled on ice. Condensed phase sample were first dissolved in 8M urea before adding loading buffer. Equal amounts of RNA were resolved on precast 15% Novex TBE–Urea gels (Thermo Fisher) in 1× TBE at ~200 V until the dye front approached the bottom. Gels were rinsed in water and stained with SYBR Gold nucleic acid stain, then imaged on a UV transilluminator. Synthetic 12- and 21-nt RNAs were run in parallel as size markers.

## Results and Discussion

### FUS LC-RGG interacts with TERRA in solution and perturbs RNA structure

To investigate how the disordered domains of FUS interact with RNA G-quadruplexes, we focused on a reduced model containing the low-complexity domain and the first RGG region of FUS (LC-RGG1, residues 1-284). We recently demonstrated that FUS LC-RGG1 recapitulates the main features of the phase separation behavior of full-length FUS (27, 28). Additionally, FUS RGG1 has been shown to specifically recognize rG4 structures (29–31). We performed NMR titration experiments by adding either TERRA12 or a single-stranded control RNA (polyU) to ^15^N-labeled FUS LC-RGG1. RNA was added in excess to suppress phase separation of FUS and keep samples in re-entrant dispersed solution regime (32), which enables monitoring of chemical shift perturbations (CSPs) in 2D ^1^H-^15^N HSQC spectra (Figure 1A). CSPs upon RNA addition were primarily observed in the RGG1 domain, indicating that both TERRA and polyU interact with this region. However, TERRA induced a distinct CSP pattern compared to polyU (Figure 1B-C), consistent with a binding mode different from non-specific association.

**Figure 1.**
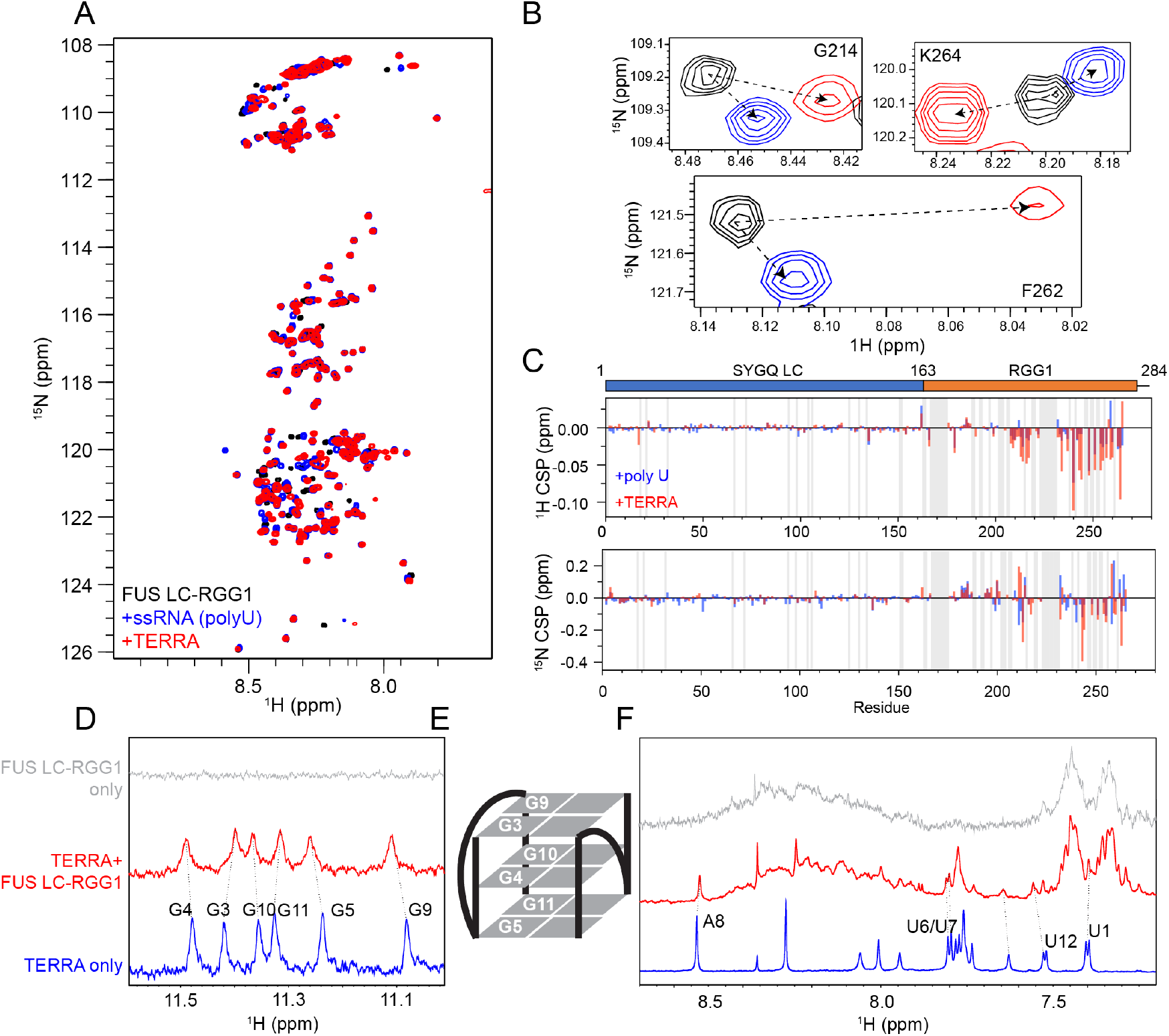
FUS LC-RGG1 interacts with TERRA in solution. A. Overlay of ^1^H-^15^N HSQC spectra of FUS LC-RGG1 (10 μM) in the absence of RNA (black), with polyU (blue, 0.43 mg/ml RNA, 25 μM protein), and with TERRA (red, 0.43 mg/ml equal to 110 μM RNA, 25 μM protein). B. Representative residues showing distinct chemical shift changes upon binding to polyU versus TERRA, indicating differential binding modes. C. ^1^H and ^15^N CSPs of FUS LC-RGG1 upon binding to TERRA (red) and polyU (blue), mapped as bar graphs by residue number. D. 1D ^1^H NMR spectra showing the imino proton region of TERRA alone (blue) and with FUS LC-RGG1 (red). E. Schematic illustration of the G-quadruplex structure formed by TERRA RNA. F. Same as D, but showing the aromatic and H8/H6 proton region (7–9 ppm), reflecting perturbations to these protons upon protein binding.

In NMR spectra of RNA, we monitored the 1D ^1^H spectra of TERRA12 RNA upon addition of FUS LC-RGG1. In the absence of FUS LC-RGG1, TERRA12 is folded with no evidence for the presence of unfolded RNA, consistent with prior literature on the high stability of TERRA12 as a dimeric rG4 structure. The FUS-rG4 interaction induce perturbation to all imino proton signals corresponding to guanine residues in the G-quartets of TERRA (Figure 1D). These include not only the top (G3, G9) and bottom (G5, G11) layers, but also the central guanines (G4, G10) (Figure 1E). As the central imino protons are buried within the core of the G4 and not directly exposed for FUS binding, the observed chemical shift changes likely reflect structural perturbations to the hydrogen bonding network that stabilizes all G-quartet stacks. Additional perturbations were observed in the aromatic region (7.2-8.9 ppm, Figure 1F), including residues in the loop region (U6, U7, A8) and the flanking ends (U1, U12), which have also been shown to contribute to rG4 stability (33, 34). We also observed peak broadening consistent with dynamic exchange between bound and unbound forms on the μs to ms timescale. Together, these results support the notion that FUS binding in dilute solution affects, but preserves, the global fold of the TERRA G-quadruplex.

### TERRA RNA is unfolded in FUS condensed phase

To assess the structure of TERRA in FUS condensed phase, we prepared macroscopic condensates of FUS LC-RGG1 as previously described (27) without and with TERRA RNA (see Supporting Information). While we were able to detect RNA resonances in the 5.6–5.9 ppm region corresponding to H1’ and H5 protons, indicating successful incorporation of RNA into the phase, the 1D ^1^H spectra revealed a complete absence of imino proton signals (Figure 2A). Imino resonances in the 11–12 ppm region are characteristic of stable hydrogen-bonded guanine pairs and are a hallmark of intact rG4 structures. The lack of these signals in the condensed phase suggests either unfolding of the rG4 structure and/or that the folded species have become NMR-invisible due to line broadening caused by restricted rotational diffusion in the viscous condensed environment.

**Figure 2.**
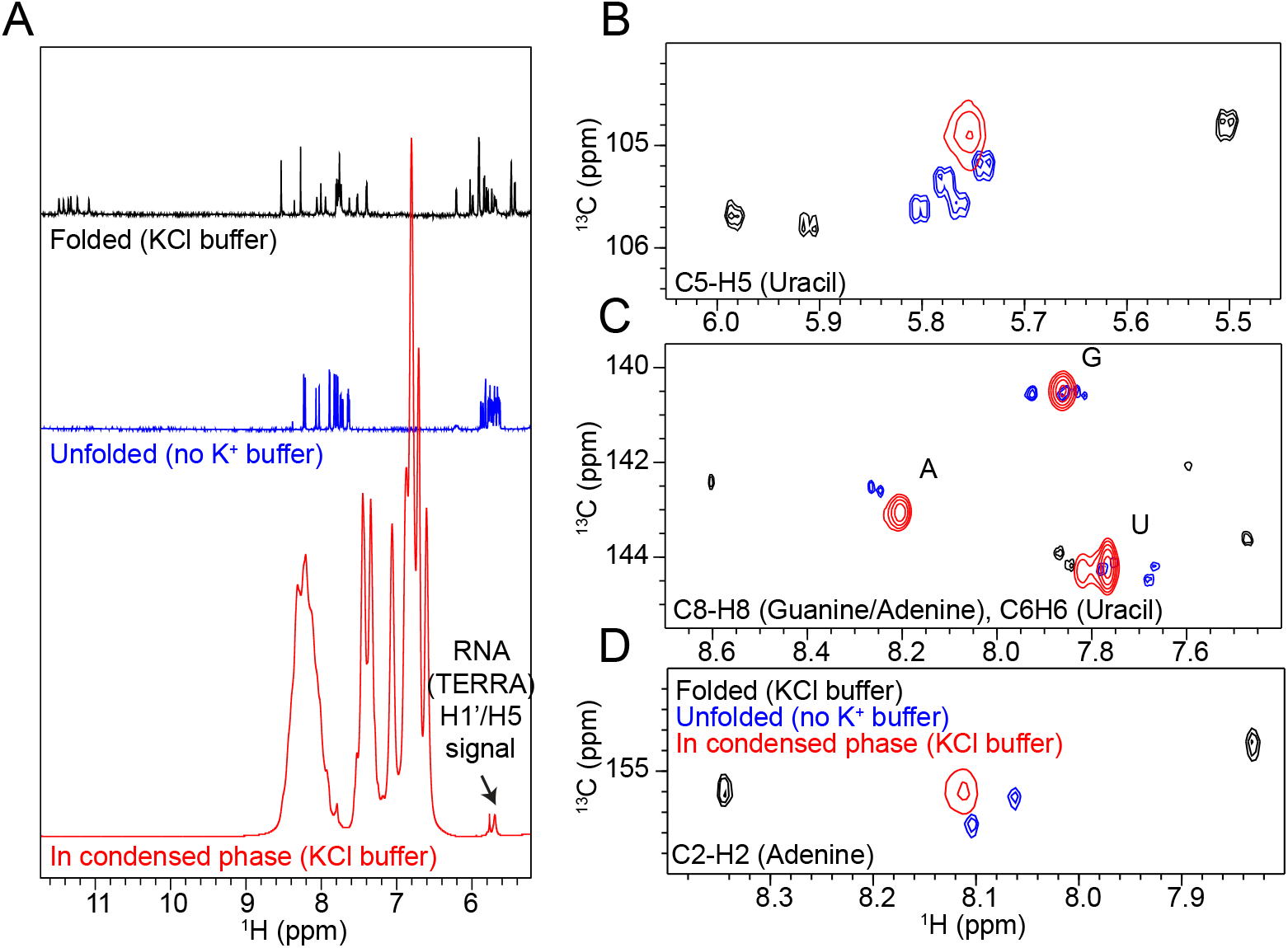
TERRA unfolds in the FUS LC-RGG1 condensed phase. A. 1D ^1^H NMR spectra of TERRA in folded (black, with KCl), unfolded (blue, no KCl), and FUS LC-RGG1 condensed phase (red) conditions. Absence of imino proton peaks and presence of a signal around 5.8 ppm in the condensed phase indicate loss of G-quadruplex structure and retention of RNA mobility. B–D. 2D ^1^H-^13^C HSQC spectra of TERRA in solution and in the FUS LC-RGG1 condensed phase. Correlations from uracil C5–H5 (B), aromatic protons C8–H8 (guanine/adenine), C6–H6 (uracil) (C), and adenine C2–H2 (D) are shown. In all cases, the TERRA signals in the condensed phase (red) overlap with those of the unfolded state (blue), indicating that TERRA is unfolded within the FUS condensed phase.

To test whether TERRA is unfolded in the condensed phase, we collected 2D ^1^H-^13^C HSQC spectra of the RNA (at natural isotopic abundance) in the condensed phase and compared them to spectra of folded TERRA in K^+^-containing buffer and unfolded TERRA in the absence of monovalent cations (see Supporting Information). Importantly, the chemical shifts observed in the condensed phase overlapped closely with those of the unfolded TERRA12 spectrum, including signals from uracil C5-H5 and C6-H6, adenine C8-H8, C6-H6 and H2-C2, and guanine C8-H8 correlations (Figure 2B-D). These resonances represent nucleotides located both in the loop and flanking regions (uridines and adenines), as well as in the G-quartet core (guanines), indicating that the observable RNA population adopts an unfolded conformation within the condensed phase. To exclude degradation as the source of these unfolded-like resonances, we assessed RNA integrity by denaturing urea-PAGE of condensed phase sample taken after NMR acquisition; a single band at the expected size with no degradation products confirmed that TERRA remained intact (Supp Fig. S1).

While these observations do not rule out the existence of NMR-invisible G-quadruplex species, they strongly support the presence of an unfolded RNA population in the condensed phase. The TERRA rG4 is a particularly robust G-quadruplex structure, with reports of >99% of the molecules folded at 25 °C (35, 36), with melting temperature in solution above 60 °C. In our hands, no unfolded species were detectable in solution by NMR under standard buffer conditions, even at elevated temperatures up to 45 ° (data not shown).

### Unfolded TERRA represents a sizable population in the condensed phase

To further support our hypothesis that TERRA adopts an unfolded conformation within FUS condensates, we employed a TERRA mutant (TERRA-mut, sequence UACCGUUACCGU) that is incapable of forming G-quadruplex structures (Figure 3A) (37). Inability of this sequence to form rG4 was confirmed by the absence of imino proton signals in K^+^-containing buffer (Figure 3B), indicating the lack of G-quartet formation. In FUS condensed phase, ^1^H-^13^C HSQC spectra revealed strong similarity between TERRA-mut and wild-type TERRA (TERRA-wt) (Figure 3C-E). These similarities included overlapping resonances from uracil, adenine, and guanine. Notably, the chemical shifts closely resemble those observed for TERRA-mut in solution (Supp Figure 2), further supporting the notion that both TERRA-wt and TERRA-mut adopt unfolded conformations within the condensate environment.

**Figure 3.**
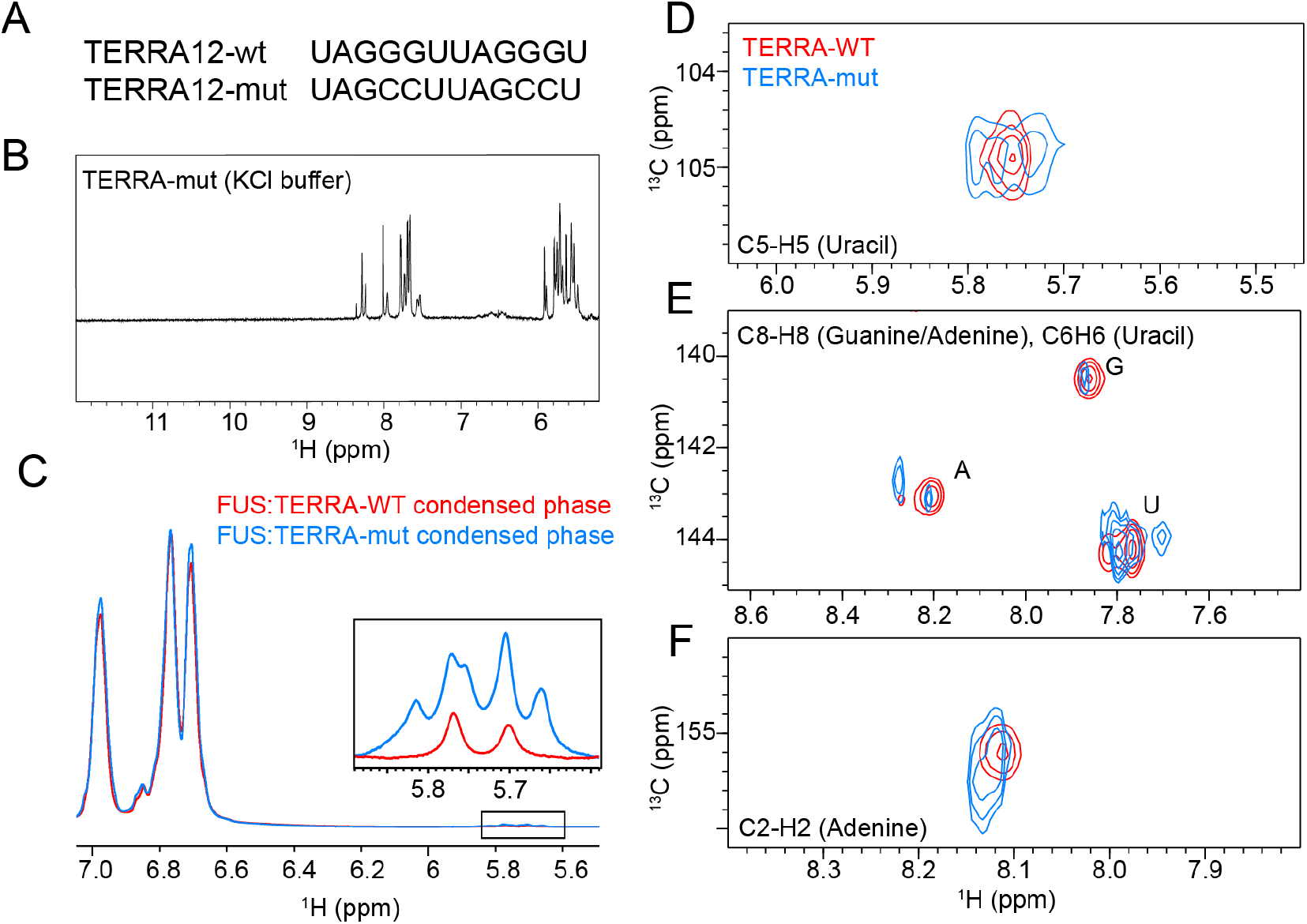
TERRA unfolding in FUS condensed phase is partial. A. RNA sequences of wild-type TERRA (TERRA-wt) and a variant (TERRA-mut) that disrupts G-quadruplex (rG4) formation. B. 1D ^1^H NMR spectrum of TERRA-mut in KCl-containing buffer shows absence of imino proton peaks, indicating inability to form a stable rG4 structure. C. 1D ^1^H NMR spectral region of protein tyrosine side-chain proton and RNA H1’/H5 resonances from FUS condensed-phase samples containing either WT or mutant TERRA. D-F. 2D ^1^H-^13^C HSQC spectra comparing TERRA-wt and TERRA-mut in the FUS LC-RGG1 condensed phase. Shown are correlations from uracil C5–H5 (C), aromatic protons C8–H8 (guanine/adenine) and C6–H6 (uracil) (D), and adenine C2–H2 (E). TERRA-mut shows similar chemical shifts but consistently higher peak intensities compared to TERRA-wt, despite equal RNA input. This suggests that only a subset of the TERRA-wt population is fully unfolded and detectable, indicating partial unfolding in the condensed phase.

To determine whether folded TERRA species might be present but not observable by NMR, and to quantify the extent of unfolding if so, we compared signal intensities of TERRA-wt relative to TERRA-mut in the condensed phase. With equal RNA input and comparable protein sidechain ^1^H signal intensities at 6.6-7.0 ppm, indicating similar condensed phase density, the total 1D ^1^H intensity of TERRA-wt was markedly lower (Fig. 3C). The same pattern held for most ^1^H-^13^C HSQC correlations including U C5-H5 and C6-H6, and A C8-H8, C6-H6, and H2-C2, with the expected exception of G C8–H8, which is weaker in the mutant because four guanines are absent per molecule (Fig. 3D-F).

Because the two RNAs are of similar length and nucleotide composition, their relaxation properties are expected to be comparable. Although chemical exchange could in principle contribute to signal attenuation, such effects would likely impact both RNAs similarly under the same conditions. Moreover, prior studies have shown that the folding process of rG4s occurs on a NMR slow timescale, and thus should not cause additional peak broadening or decrease signal intensity for the unfolded species (38). Therefore, peak intensity serves as a reasonable proxy for estimating the relative population of unfolded RNA. Based on the comparison of RNA H1’ and H5 ^1^H peak intensities, we estimate that approximately 30-40% of TERRA-wt is unfolded and detectable by NMR in the FUS condensed phase, while the remaining 60-70% likely exists in a folded state that is broadened beyond detection due to restricted dynamics or conformational exchange within the viscous, condensed environment. The markedly reduced peak intensity of TERRA-wt relative to TERRA-mut thus supports the interpretation that at least a sizable population of the total TERRA-wt population is unfolded and NMR-visible in the condensed phase.

Taken together, we show that even the unusually strong rG4 structure of TERRA is destabilized in biomolecular condensates (Figure 4). Specifically, entry into the FUS condensed phase leads to unfolding of at least a substantial fraction of TERRA RNA. These data may help resolve the controversy surrounding the apparent absence of rG4 structures in cells (9) As condensates containing RGG sequence regions can unfold rG4s, the RNA may transiently become susceptible to chemical labeling and are subsequently interpreted as non-folded. Condensate-driven rG4 unfolding could also be biologically functional, representing a regulatory mechanism for RNAs whose activity depends on their structural state, such as those involved in telomere maintenance, translational repression, or repeat-associated toxicity. The ability may be due to the unique environment of the condensate where the dielectric constant disfavors other structures like RNA duplexes (39) or simply due to the high concentration of RGG motifs that bind rG4s.

**Figure 4.**
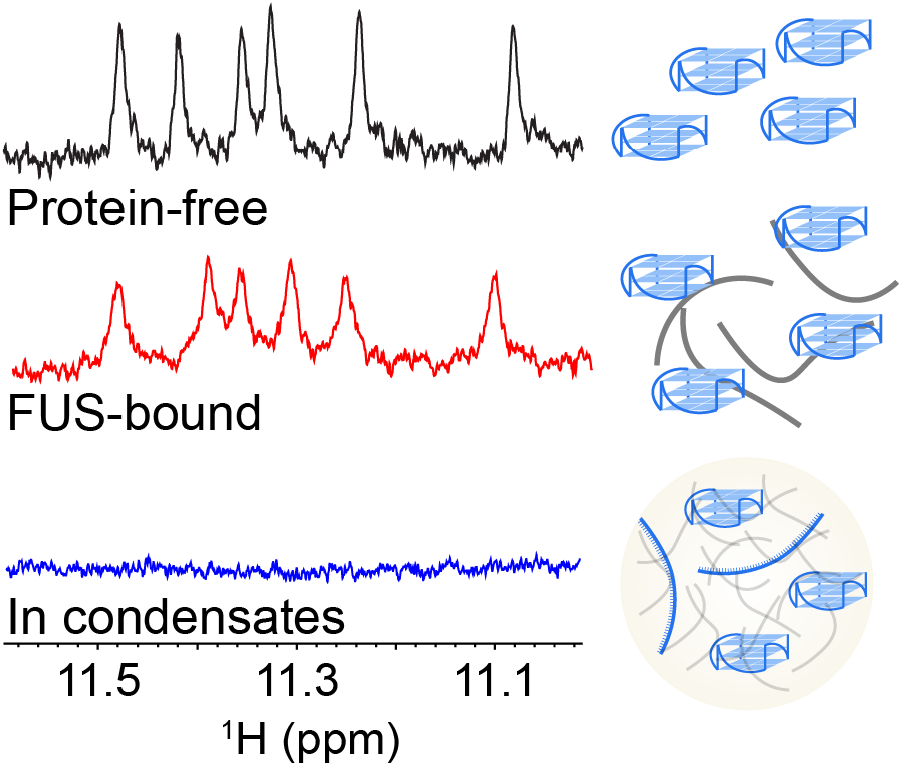
Summary Model of TERRA and FUS interaction in condensation. TERRA RNA structure is perturbed by FUS in the dispersed phase but significantly disrupted including at least a sizeable fraction that is completely unfolded in FUS condensates.

More broadly, our work illustrates how the physical/chemical properties of condensed phases, in conjunction with protein-RNA interactions, can reshape RNA structure and influence its fate, accessibility, or function in the cellular environment. These observations support a model in which biomolecular condensates act not only as organizational platforms, but also as dynamic regulators of RNA conformation and function.

## CRediT authorship contribution statement

**Tongyin Zheng:** Writing – review & editing, Writing – original draft, Methodology, Funding acquisition, Formal analysis, Conceptualization.

**Nicolas L Fawzi:** Writing – review & editing, Funding acquisition, Conceptualization.

## Acknowledgements

The authors would like to thank Dr. Mandar Naik (Brown University) for assistance with NMR and the Structural Biology Core Facility at Brown University. Research was supported in part by NIGMS R01GM147677 (to NLF). TZ was supported in part by a Pape Adams Postdoctoral Award from the Carney Institute for Brain Science at Brown University and a Milton Safenowitz Postdoctoral Fellowship (23-PDF-629) from the ALS Association. This content solely reflects the authors and does not necessarily represent the official views of the funding agencies.

