## Supplementary Figures for "Visualizing TERRA RNA G-quadruplex unfolding in FUS biomolecular condensates"

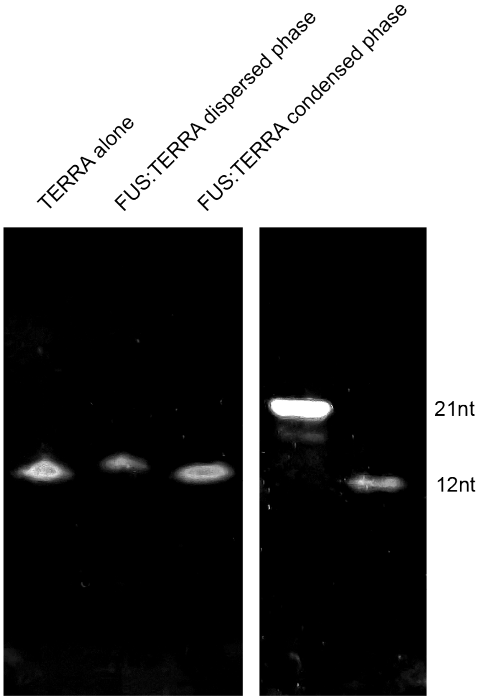


**Supp Figure 1. TERRA remain intact in FUS condensed phase.** Denaturing urea-PAGE shows that TERRA RNA from urea-dissolved FUS condensates migrates as a single band at the expected length, indicating no detectable degradation. RNAs of known length are displayed as standards on the right.

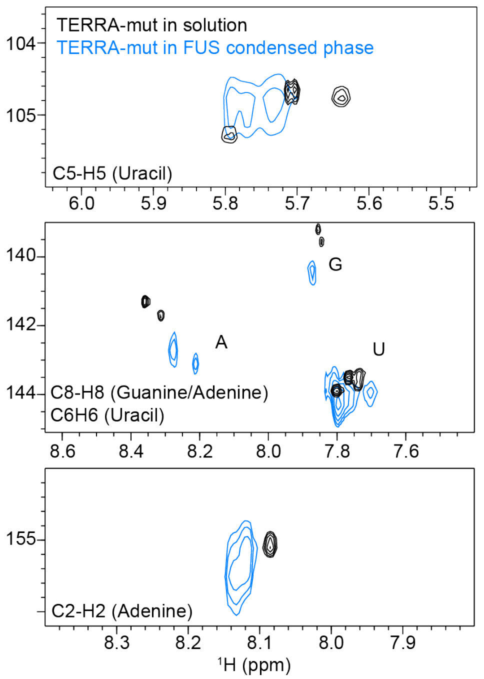


**Supp Figure 2. Chemical shifts of TERRA-mut in the condensed phase closely resemble those observed in solution.** 2D NMR spectra ($^{15}$N–$^1$H HSQC) of TERRA-mut in solution (black) and in the condensed phase (blue) demonstrate strong overlap of peak positions, indicating similar RNA conformations in both environments.
